# An Artificial Intelligence-Assisted Digital Microfluidic System for Multistate Droplet Control

**DOI:** 10.1101/2023.11.09.566344

**Authors:** Kun-Lun Guo, Ze-Rui Song, Jia-Le Zhou, Bin Shen, Bing-Yong Yan, Zhen Gu, Hui-Feng Wang

## Abstract

Digital microfluidics (DMF) is a versatile technique for parallel and field-programmable control of individual droplets. Given the high freedom in droplet manipulation, it is essential to establish self-adaptive and intelligent control methods for DMF systems with informed of the transient state of droplets and their interactions. However, most related studies focus on the localization and shape recognition of droplets. Here, we develop an AI-assisted DMF framework named “μDropAI” for multistate droplet control based on droplet morphology. Semantic segmentation model is integrated into our custom-designed DMF system to recognize the droplet states and their interactions for feedback control with a state machine. The proposed model has a strong generalization ability and can recognize droplets of different colors and shapes with an error rate of less than 0.63%. It enables control of droplets without user intervene. The proposed system will inspire the development of semantic-driven DMF systems which can interface with artificial general intelligence (AGl) models for fully automatic control.

## 1 Introduction

Digital microfluidics (DMF) has become a prevalent technique for handling liquid due to its unique flexible and discrete fluid control capabilities.^1–3^ In particular, DMF systems based on electrowetting-on-dielectric (EWOD) principles utilize voltage modulation to control the wettability of solid surfaces, facilitating the manipulation of droplets across scales from nL to μL.^4–7^ This technique has advantages such as miniaturization, programmability, parallelization, and low power consumption.^8, 9^ Due to these benefits, DMF devices have been used in a wide range of applications, including point-of-care testing,^10, 11^ cell research,^12–14^ biomedicine,^15^ and environmental monitoring.^16^

Typically, DMF devices accept a standard set of basic instructions to perform droplet manipulations. Users can create droplet actuation sequences for each actuating electrode, ^17, 18^ thereby allowing the droplets to move, dispense, merge, split, and mix. ^19–21^ Typically, an automated DMF system incorporates sensors liking capacitance or impedance to monitor droplet operations in real-time.^22–25^ The control algorithm is also integrated into the DMF system to manage electrode switching and sequencing for subsequent operations, including droplet tracking and path planning based on feedback signals.^26, 27^ These sensing and control methods not only enhance the droplet manipulation stability but also optimize the motion performance of the droplet.^26^ Imaging technologies, which include background subtraction methods, edge detection, and image segmentation,^28–30^ are applied to droplet motion recognition. They offer a non-contact, real-time solution for signal acquisition, resulting in rapid and accurate detection and tracking of droplets, even transparent ones. ^31^ In practice, the operations of droplets often result in changes of droplet appearance such as shapes and colors. However, few studies have focused on the automated control of droplets with recognition of their states based on the appearance changes.

In the last decade, the advance of convolutional neural network paved the way for the development of semantic segmentation techniques, which is a pixel level classification that provides the understanding of images.^32^ Compared to the image processing such as edge detection and YOLO,^33^ the semantic segmentation techniques stands out for its outstanding advantages in complex scenarios and multi-category object segmentation tasks, including precise object segmentation, support for multiple categories, environmental awareness, and advanced image analysis applications.^34–36^ Therefore, they have been widely applied in AI-assisted technology, such as autonomous driving, robotics, and smart healthcare. ^37–39^ Therefore, semantic segmentation techniques can provide DMFs with accurate recognition of droplet states, complemented by control methods that enable the development of fully automated and flexible control systems to perform complex tasks.

In this paper, we develop an artificial intelligence-assisted digital microfluidics system named “μDropAI” for highly automated multistate droplet control (Fig. 1). The framework contains four parts: 1) a hardware system for droplet actuation and video capture; 2) a semantic segmentation model trained to detect the droplet states; 3) a region growth algorithm to extract the position and morphological information of droplets; and 4) automated control processes for user-programmed automatic control. An open-source dataset was established with labelled droplet images consisting different states, colors and shapes. The proposed DMF system had a semantic segmentation model trained on the dataset to recognize droplet multistate, controlled the electrode energization sequence, and was successfully used for automatic droplet segmentation, movement, and dispensing. To our best knowledge, it is the first time to use the semantic segmentation for the control of DMF systems. Both the dataset and the μDropAI are open source. In addition, the proposed control system is also compatible with existing DMF devices that integrated with digital camera such as DropBot and openDrop.^40, 41^ Therefore, it can be widely used in DMF-based applications and it will promote the integration of more AI-based techniques to be utilized in DMFs such as reinforcement learning and ChatGPT.^42^

**Fig. 1.**
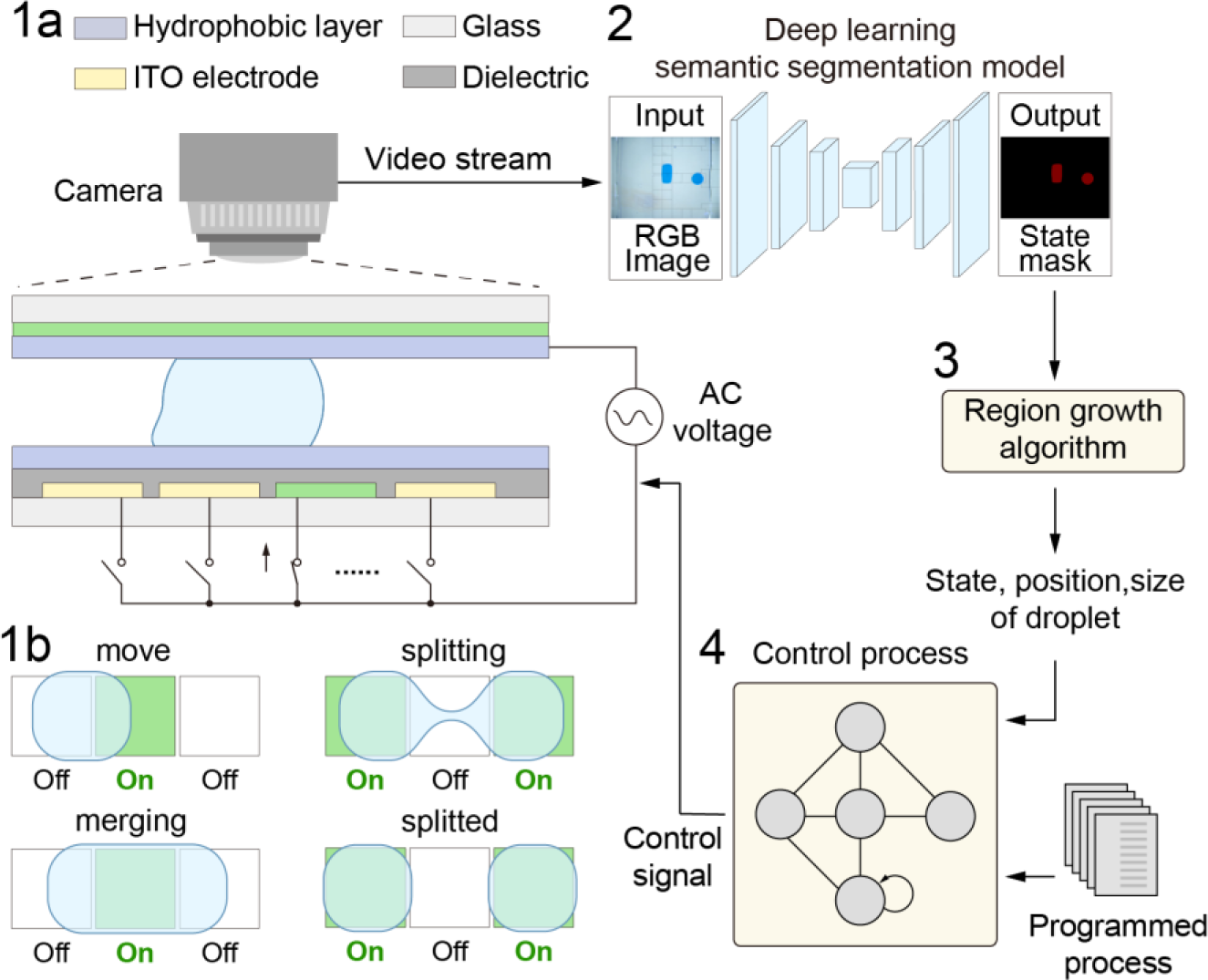
System overview of the artificial intelligence-assisted digital microfluidic system named “μDropAI”. From left to right, from top to bottom, in sequence: 1a) a hardware system for droplet actuation and video capture; 1b) droplet control effect; 2) a semantic segmentation model trained to recognize the droplet states at the pixel level; 3) a region growth algorithm to extract the position and morphological information of droplets; 4) automated control processes for user-programmed automatic control.

## 2 Methodology

### 2.1 Encoder-decoder semantic segmentation model

A deep learning-based algorithm which automatically segments droplet multistate is proposed. Our model is based on the U-net model with an encoder-decoder structure.^43^ The encoder unit of the proposed model performs feature extraction and the decoder unit performs upsampling operations. Shortcut connections connect the corresponding feature maps between the encoder and decoder units (Fig.2). By incorporating skip connections, the model can capture both high-level and low-level features.

**Fig. 2.**
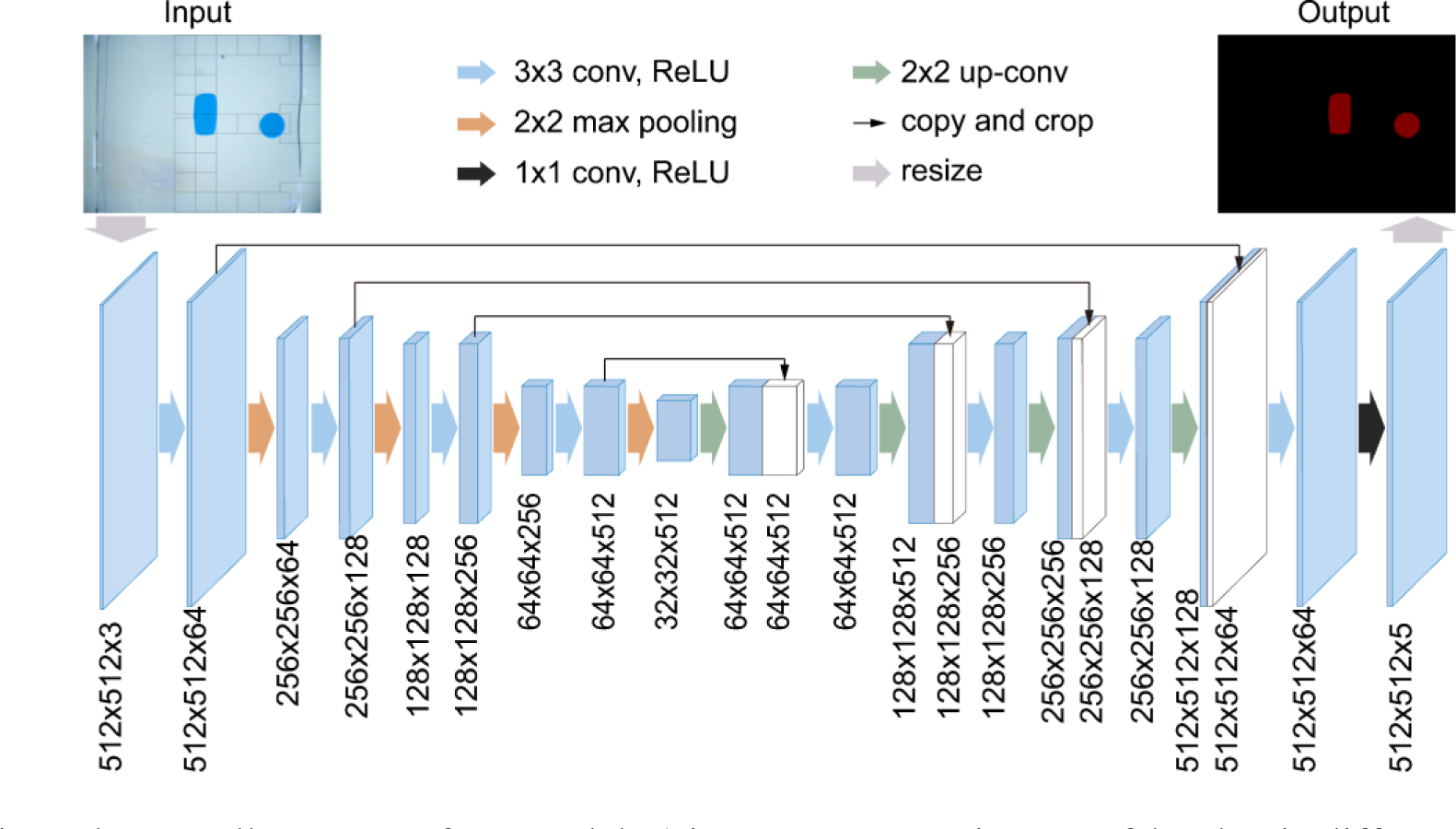
The overall structure of our model. 1) input 1920x1080 images of droplets in different states; 2) downsampling of image pixel and channel count (512x512 → 256x256, 64 → 5), and upsampling of image pixel and channel count (256x256 → 512x512, 3 → 64); 3) output 1920x1080 labelled images.

In the encoder stage, the model comprises 16 convolutional layers and 3 fully connected layers to extract crucial feature information from input images, which is later used in the subsequent decoder stage to generate semantic segmentation masks. These convolution layers all use a 3x3 convolution kernel, a step length is 1, and the same filling method to keep the size of the feature graph unchanged. After each convolution layer, the ReLU activation function is applied to introduce nonlinearity.

The convolution layer part can be divided into 5 convolution blocks. There are 2 convolution layers within each convolution block, followed by a 2x2 maximum pooling layer, which is used to reduce the spatial size of the feature graph. By stacking multiple convolutional blocks, the depth of the network can be gradually increased, thus improving the feature expression ability of the model.

In the decoder stage, 8 convolutional layers corresponding to the encoder are used, followed by upsampling and transpose convolution operations after each block. These operations restore the feature map size to the original input image size, and skip connections are utilized to connect the feature maps of the same resolution from the encoder and decoder stages. These shortcut connections help retain more spatial information and details, thus assisting the network in obtaining more accurate segmentation results.

The final layer of the network is a 1x1 convolutional layer with 5 channels. This layer maps the feature maps from the decoder to the same number of channels as the number of label categories considered in this paper. This approach ensures that the predicted image segmentation results are accurate.

### 2.2 Region growing algorithm

The region growing algorithm (†ESI S1 and †Fig. 1) is introduced to obtain a pixel-level division of the droplet state in each frame. Additionally, these segments can be connected based on pixel similarity to obtain a more accurate representation of the droplet. This algorithm has a strong ability to segment and compute regions based on pixel similarity.^44^

### 2.3 Automated control process based on droplet states

The control process is executed using a state machine implementation, enabling automated manipulations, such as droplet movement and splitting, following user-provided manipulation instructions (Fig. 3). For example, by inputting the move process steps, the droplet moves in the direction of the specified electrode, and the control method determines whether the droplet has reached the specified position according to the feedback information (Fig. 3a). When the split command is input, the droplet is controlled to split. During the splitting process, the droplet states are recognized, and the next manipulation is determined based on the feedback information to ensure that the droplet has completely split (Fig. 3b). When the initial state contains two droplets, a merge command can be applied to merge the two droplets into one large droplet (Fig. 3c).

**Fig. 3.**
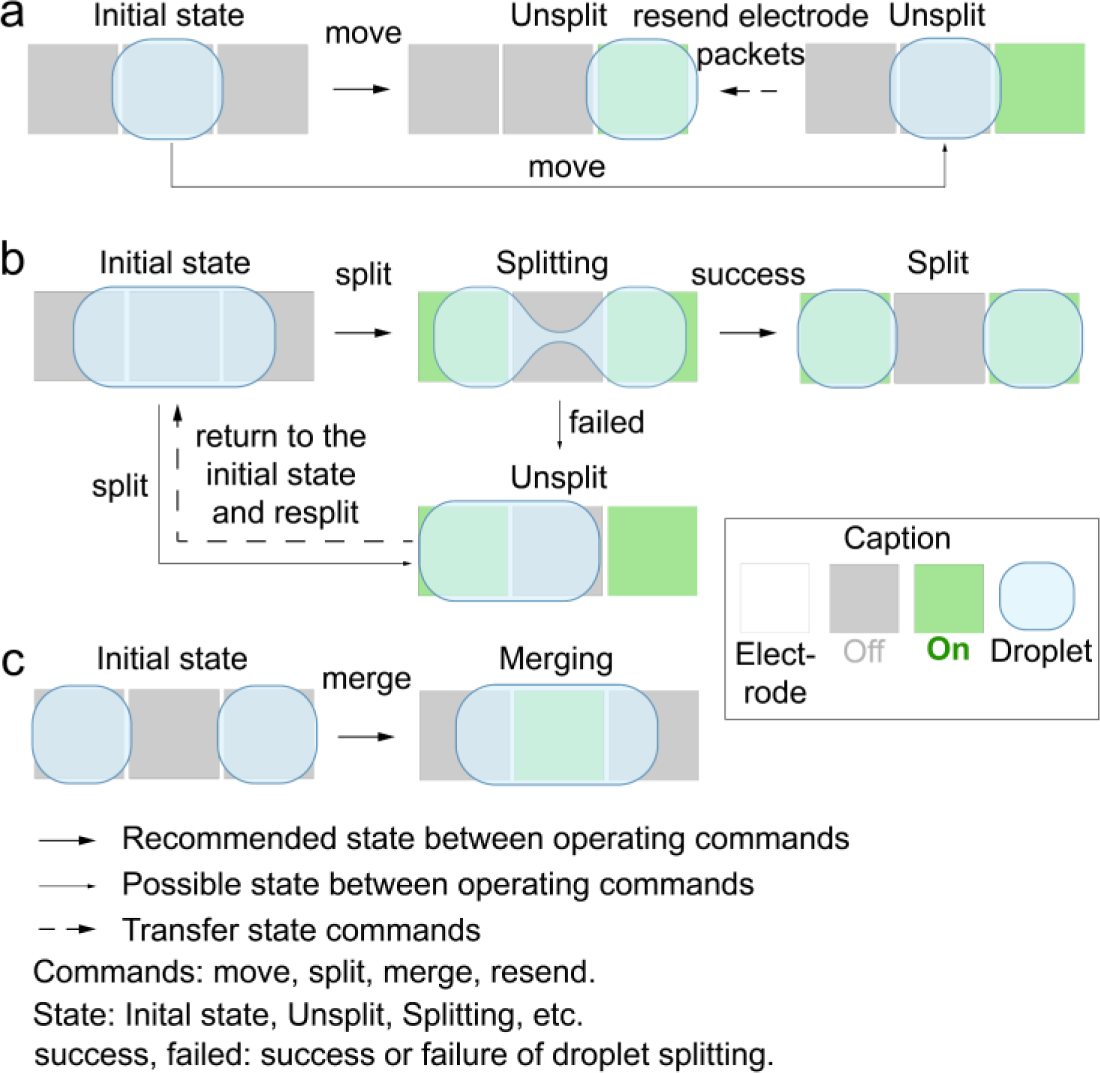
Schematic diagram of the droplet states in the control processes. (a) Droplet movement control process based on states; (b) Droplet splitting and movement control process based on states; (c) Droplet merging control process based on states.

## 3 Experimental methods

### 3.1 Chip fabrication

A double-plate DMF chip was utilized, with both the top and bottom plates constructed from ITO glass. The electrodes were directly etched onto the bottom plate (52 mm x 50 mm) using a UV laser operating at a wavelength of 355 nm and a pulse duration of 20 μs. ^45^ Then, the glass plates were cleaned with ethanol for 10 minutes to remove residual material and exposed to plasma treatment for 1 minute. Following the application of a vacuum-deposited Parylene C film (∼2 µm thick) onto the bottom plate surface as a dielectric layer, CYTOP films were spin-coated (2000rpm, 30s) onto both the top and bottom plates to serve as hydrophobic layers. A double-sided adhesive with a thickness of 0.5 mm was employed to securely attach the upper and lower plates. Then, silicon oil (0.65 cs) was introduced to fill the gap between the two plates, effectively diminishing the friction encountered during liquid droplet movement and deterring evaporation.

### 3.2 Setup of DMF hardware

A homemade DMF system was used to manipulate the DMF chip and control the droplet manipulation processes. Optocoupler switch array controlling the high voltage (ATG-2081, Aigtek, China) applied to the electrodes on the chip was configured by a microcontroller unit (STM32L432, ST, Italy), which receives the instructions from a python script embedded with the proposed AI assisted control model. A microscope equipped with a CCD camera (IUA20000KPA, ToupTek, China) was positioned above the DMF system to observe and capture the droplet status. The resolution of the CCD camera was set to 2560x1440 to obtain clear images. Subsequently, the images were transmitted to a computer with an R7 5700X CPU, 16 GB RAM, and RTX 3070 GPU for further processing and analysis.

### 3.3 Dataset establishment

A multistate droplet dataset was established with a digital camera. Images of various droplet manipulations were captured with frame rate of 30 frames per second. The captured videos were converted into a series of images (1920x1080) with OpenCV. These droplet manipulation images were labelled into four types of states: “unsplit”, “splitting”, “split”, and “merging” with LabelMe (†ESI Table 1), and converted to PASCAL VOC data format. The dataset was divided into training and test sets in a ratio of 0.8:0.2, including 1751 images in the training set and 438 images in the test set. The training set was used to train the model, and the test set was used to evaluate the model’s performance. The dataset had been made public for others (https://github.com/Eric1105/-DropAI.git). This protocol can be used as a reference for establishing other DMF droplet manipulation datasets.

### 3.4 Model training parameters and evaluation metrics

The training GPUs were dual RTX3090 12G. The experiment was conducted using the PyTorch framework, with the programming language Python version 3.8 on a Windows system.

To adapt to the dataset, which has a large proportion of background and a small proportion of objects, the following optimized parameter was set to train the semantic segmentation model. After our testing, Adam was chosen as the optimizer it has the ability to quickly identify the optimal parameter configuration to achieve better training performance. The learning rate was set to 0.0001, with a minimum learning rate of 0.000001. The learning rate was reduced every 5 epochs using the cosine annealing learning rate scheduling strategy.

The Dice loss was employed as the loss function (Eq.1). The Dice loss ranges from 0 to 1, with lower values indicating greater similarity between the predicted and ground truth results.

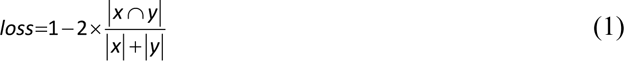

where *x* represents the predicted segmentation mask and *y* represents the ground truth segmentation mask.

The mean intersection over union (mIoU) was employed as the main evaluation indicator for the semantic segmentation performance (Eq.2). This metric is the average of the ratio of the intersection and union of various real labels and prediction results.

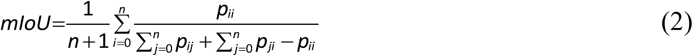

where *n* represents the total number of categories, *p_ii_* represents the number of correct predictions, *pij* represents false negatives, in which *i* is predicted as *j*, and *p_ji_* represents false positives.

## 4 Results and discussion

### 4.1 Model comparison

To validate the segmentation performance of the proposed deep learning algorithm, a comparative experiment with the traditional segmentation model U-net and DeeplabV3+^46^ on our open-source dataset was designed to compare their segmentation results, mIoU, and loss. The mIoU indicates the extent to which the model-predicted area overlaps with the actual area. The loss is used to measure the difference between the model prediction and the actual target. In the recognition process, red represents the “unsplit” state, green indicates the “splitting” state, yellow represents the “split” state, and blue signifies the “merging” state.

Fig. 4a-d illustrates the process of a droplet splitting. As can be seen, it transitions from the initial red “unsplit” state to the green “splitting” state, culminating in the yellow “split” state. In our evaluation, the DeeplabV3+, U-net, and our proposed model accurately recognize droplets in the “unsplit” and “split” states (Fig. 4b1-d1 and b3-d3). In instances where droplets assume an hourglass shape, the DeeplabV3+ fails to segment the connected regions within the “splitting” state (Fig. 4b2). In contrast, both U-net and our proposed model effectively segment the “splitting” droplet (Fig. 4c2 and d2). Accurate recognition of the “splitting” state is crucial for the successful execution of the splitting operation. When two droplets merge, our proposed model accurately discriminates between the blue “merging” and “split” states (Fig. 4d4). In contrast, the DeeplabV3+ erroneously categorizes them as “split” (Fig. 4b4), while U-net detects parts of the droplets as “split” (Fig. 4c4). These results indicate that our model consistently outperforms the other models in most scenarios.

**Fig. 4.**
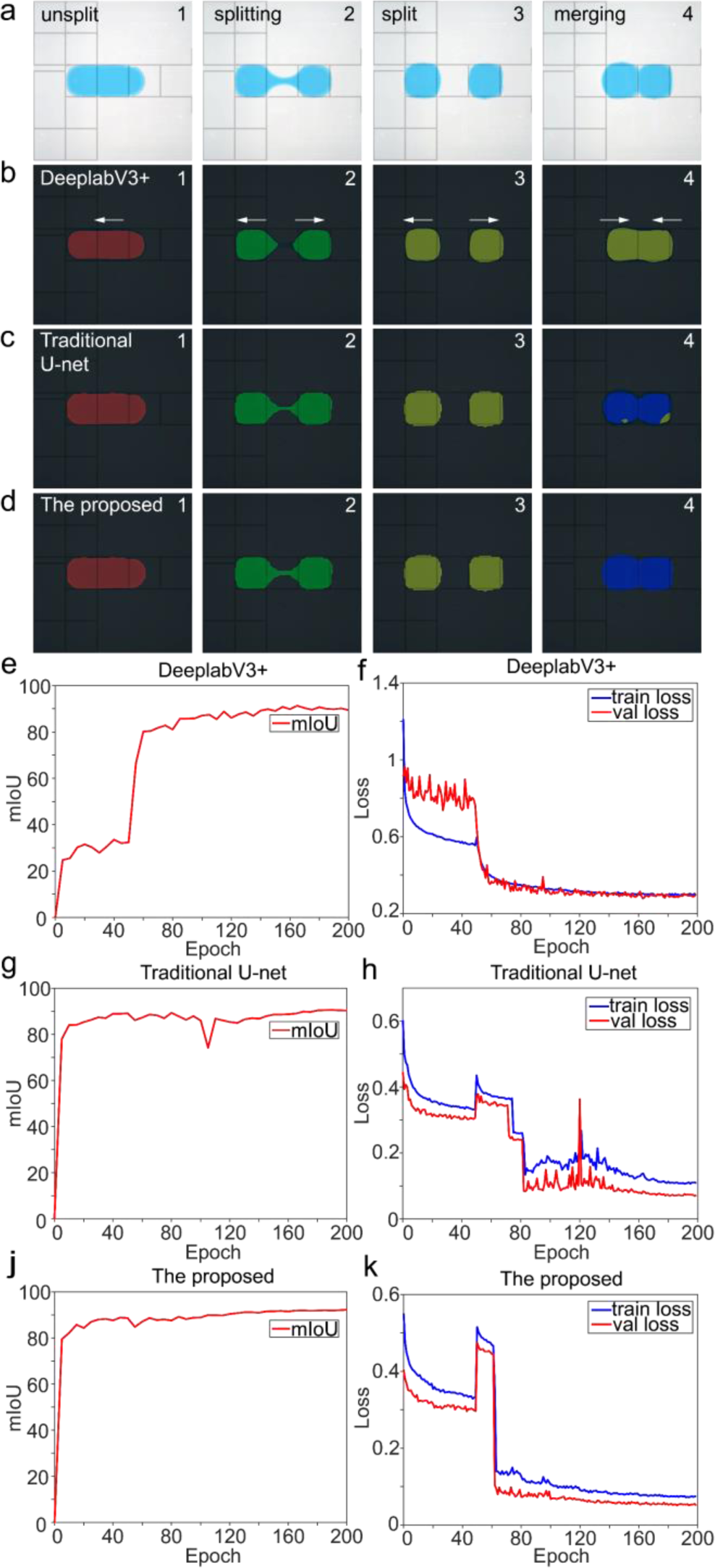
Training and evaluation of the semantic segmentation model. (a) Original images; (b) Segmentation results using the DeeplabV3+; (c) Segmentation results using the traditional U-net; (d) Segmentation results using the proposed; (e) ThemIoU of the DeeplabV3+ (The mIoU indicates the extent to which the model-predicted area overlaps with the actual area); (f) The loss of the DeeplabV3+ (The loss indicates the difference between the model prediction and the actual target); (g) The mIoU of the traditional U-net; (h) The loss of the traditional U-net; (j) The mIoU of the proposed; (k) The loss of the proposed.

The mIoU of the proposed is in the range of 85-91% (Fig. 4j), while the mIoU of DeeplabV3+ and U-net are in the range of 80-89% and 83-89% (Fig. 4e and g). The mIoU of the proposed 1.1-6% better than that of the DeeplabV3+, and 1.1-2% better than that of the U-net. A higher mIoU indicates better segmentation performance, with the proposed predicted regions overlapping more closely with the actual regions. Moreover, the loss of the proposed stabilizes at 0.08-0.15 (Fig. 4k), which is approximately 50-60% lower than the DeeplabV3+ loss of 0.2-0.3 (Fig. 4f), and 20-37% lower than the U-net loss of 0.11-0.18 after the 75th epoch (Fig. 4h). A lower loss indicates that the proposed predictions are closer to the true droplet states. Furthermore, the loss and mIoU of the proposed tend to stabilize after convergence, indicating that it has learned and mastered effective semantic segmentation ability.

The pixel-level evaluation metrics on a test set of 438 images were also computed, including the mean precision (mPrecision), mean pixel accuracy (mPA), and mean recall (mRecall) values. Three metrics are used to evaluate the performance of the model’s semantic segmentation results in terms of category accuracy, pixel-level classification accuracy, and positive sample detection (†ESI S2 Eqs.1-5). The mPrecision is 95.72%, the mPA is 93.30%, and the mRecall is 93.30% (†Fig. 2). The three metrics of the proposed method are greater than 91.76%, 89.80%, and 88.76% of the DeeplabV3+, and greater than 92.96%, 90.97% and 90.37% of the U-net, with higher values indicating that the proposed can accurately recognize objects in the image and generate segmentation results that highly match the actual labelled objects.

### 4.2 Influence of luminous environment

To investigate the impact of potential variations in luminous environment within various DMF systems, OpenCV was utilized to adjust the brightness levels of real-time videos (Fig. 5a) and the accuracy of recognizing states, positions, and overall recognition under different lighting conditions was analyzed across eight videos (†ESI S2 Eqs.6-8).

**Fig. 5.**
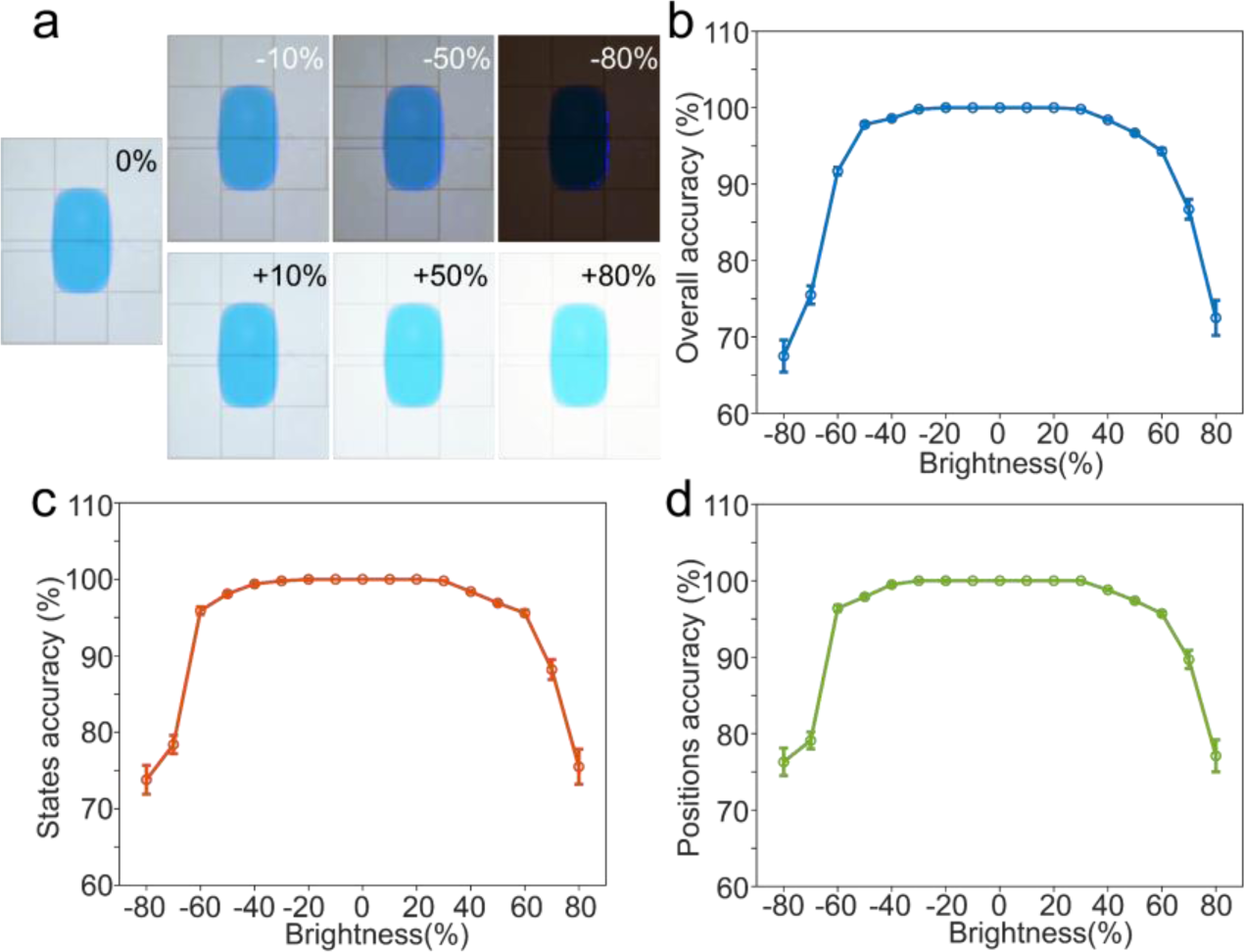
Recognition and segmentation results of 8 different real-time videos under different brightness levels. (a) Comparative images, showing the original image and images with different brightness levels; (b) Overall mean accuracy (±standard deviation, SD) of different real-time videos at various brightness levels; (c) State mean accuracy (±SD) of different real-time videos at various brightness levels; (d) Position mean accuracy (±SD) of different real-time videos at various brightness levels.

With a 0% and 50% increase or decrease in brightness, the overall accuracy decreased to 100% and 96.7% (Fig. 5b). Meanwhile, the mean accuracy for recognizing both states and positions consistently ranges between 97% and 100% (Fig. 5c and d). Obviously, the DMF’s lighting just changed the droplet color from blue to cyan or dark blue, and the droplet on the DMF chip are still visible. The change in lighting intensity is consistent with the lighting conditions of the vast majority of DMFs, and in this case, the proposed system still maintains a high recognition accuracy of over 96.7%. With a 60% and 80% increase or decrease in brightness, the overall mean accuracy decreases to 91.7% and 67.5% (Fig. 5b). Likewise, the mean accuracy for states and positions recognition decreases to 95.6% and 73.8% (Fig. 5c), 95.7% and 76.3% (Fig. 5d). It is evident that when the brightness variation approaches 80%, overexposure or underexposure occurs. The droplet’s color has changed, and its edges are no longer distinct, leading to a noticeable drop in recognition accuracy. Furthermore, as the brightness varies from 0% to 50%, the standard deviation (SD) of overall accuracy consistently remains within the range of 0-0.3%, which is lower by 0.5% to 2% compared to the observed SD fluctuating between 0.5% to 2.3% when brightness varies from 60% to 80%. This indicates that the proposed system exhibits stronger accuracy and robustness, especially in the brightness variation range of 0-50%.

### 4.3 Influence of droplet color and shape

Considering the difference in color and shape of droplets due to differences in their composition and handling methods in DMF system, experiments for recognizing droplets with different colors and shapes were established. Droplets of different colors can be recognized as “unsplit” by the different shapes they appear to take during moving, such as L-shaped, rectangular, circular, and triangular. (Fig. 6a1 and †Fig. 3a). In real-time operation, the state transitions of the splitting of different colored droplets from “unsplit” to “splitting” to “split”, and the merging of two droplets “merging” can also be recognized (Fig. 6a2-5, †Fig. 3b-e, and †video 1). In conclusion, the proposed system not only accurately recognizes various shapes and states of blue droplets but also extends its recognition to red, yellow, black, and even transparent droplets (†ESI S4). This proves that our system has strong generalization ability to recognize different colors and shapes droplets of instantaneous changes in operating states.

**Fig. 6.**
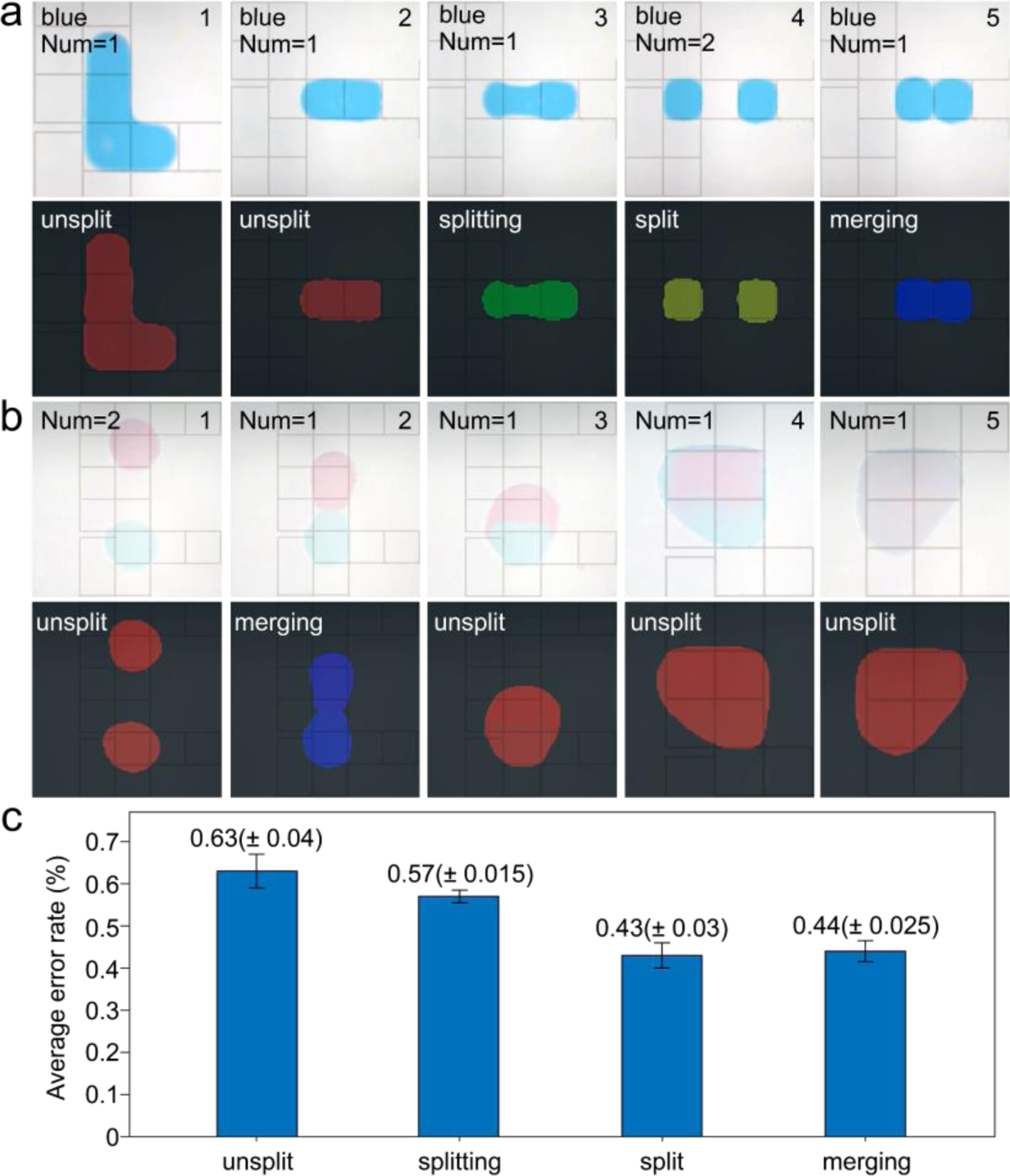
Recognition and segmentation results for droplet color and shape. (a) The droplet recognition results under different states (Num: number); (b) The merging recognition results of two different colored droplets; (c) The average error rate (±standard deviation, SD) of recognition in droplet manipulation with different states and colors (red, yellow, blue, black, and transparent). The error rate is the error caused by the droplets being recognized as other states while remaining in the continuous and same state.

Considering the merging of droplets with various substances during DMF experimental operations to observe reaction phenomena, an experiment for the merging of droplets of different colors was devised. When red and blue “unsplit” droplets merge (Fig. 6b1), they enter a “merging” state (Fig. 6b2), returning to “unsplit” state after merging (Fig. 6b3). Even as they mix further, they still stay “unsplit” and trackable (Fig. 6b4-b5)). The successful recognition of the merging of droplets with different colors indicates the ability of the proposed DMF system for merging, subsequent splitting, and moving of droplets with various reagents or materials, extending beyond the constraints of single-substance droplets. The error rate is the error caused by the droplets being recognized as other states while remaining in the continuous and same state. It can be found that the average error rates for the 4 states of different colored droplets are maintained at around 0.63%, 0.57%, 0.43% and 0.44% respectively, which means that the average number of incorrectly recognized frames is less than 1 frame in continuous and same state (Fig. 6c). This demonstrates the system’s capability to perform real-time recognition of multiple states of differently colored droplets with a low error rate. When applied in real-time recognition during DMF experimental operations, it ensures the automation of droplet handling processes, preventing operational failures due to misrecognized states. For example, in a control sequence that transitions from a “splitting” state to a “splitting” state that is incorrectly identified as “splitting,” the system incorrectly determines that the droplet has completed the splitting process.

### 4.4 AI-assisted multistate droplet control

To validate the applicability of the AI-assisted multistate feedback control system named μDropAI, droplet manipulations including moving, splitting, and dispensing on the DMF platform were performed. In Fig. 7b and c, the horizontal axis represents the nu mber of frames in the experiment, and the vertical axis represents the position of the droplets in the same frame, multiple lines are utilized to indicate the droplet’s positions in each frame.

**Fig. 7.**
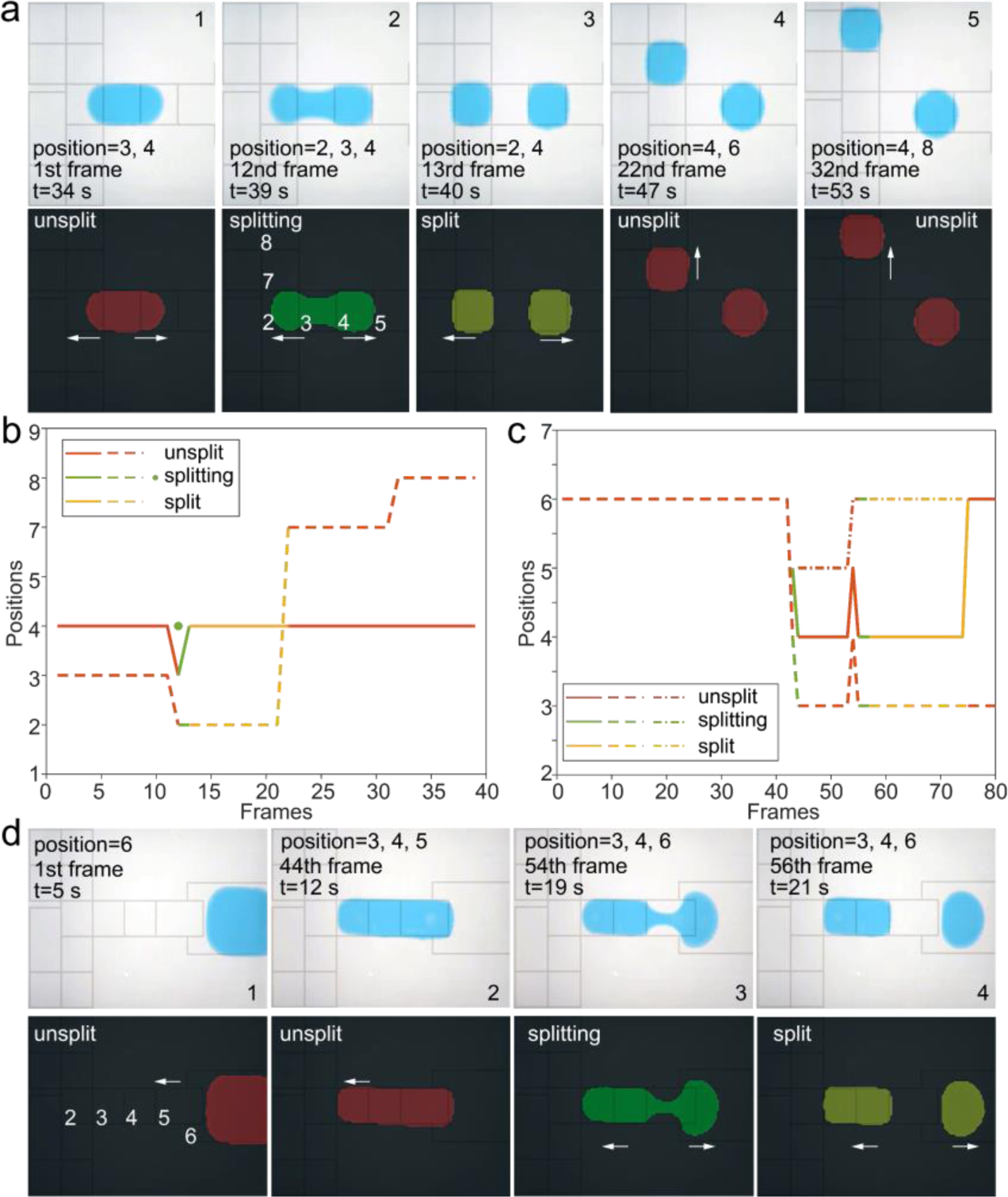
Processes and results of droplet moving, splitting, and dispensing. (a) The states and positions during droplet moving and splitting processes in different frames; (b) The state and position results in droplet moving and splitting processes; (c) The state and position results in droplet dispensing from liquid reservoir; (d) The states and positions of droplet dispensing from the liquid reservoir in different frames.

Droplet moving, and splitting on digital microfluidic are common biological and chemical experimental operations. A flow chart of automated feedback control involving droplet moving and splitting is performed (†ESI S5 and †Fig. 4). In the droplet moving manipulation, μDropAI recognizes the droplets and determines their locations on the platform. Subsequently, the activated electrodes drive the droplets to reach their destination. When the droplet reaches the target electrode, μDropAI confirmed that the droplet has successfully moved. (Fig. 7a4-a5, Fig. 7b, and †video 2). When the movement fails, the electrode needs to be reactivated to drive the droplet. Compared to existing image-based methods,28 our system not only recognizes the contours of the droplets, but also distinguishes the state and position of the droplets and is adaptable to a variety of environments. In the droplet splitting manipulation, the droplet undergoes a total of three state transitions, from the “unsplit” state of a large droplet to the hourglass-shaped “splitting” state, and finally to the “split” state of two small droplets (Fig. 7a1-a3, Fig. 7b, and †video 2). When two electrodes are energized sequentially, the droplet may be pulled to one side by the first energized electrode, leading to a failure in splitting. In such cases, the control system reverts the droplet to its initial state and re-split (†video 3). Additionally, the system can automatically merge the droplets and to recognize this droplet. The proposed control system achieves real-time monitoring of droplet states and guides droplet operations based on their states, extending beyond mere location-based operation determination.

Droplet dispensing is an operation to generate droplets from a reservoir. A flow chart of automated feedback control experiments for dispensing is performed (†ESI S5 and †Fig. 5). The initial position and state of droplet in reservoir is recognized (Fig. 7d1), an automatic process is executed by the μDropAI to activate the electrodes to split the droplet from the reservoir, (Fig. 7d2-d3). During the process, an hourglass-shaped structure is formed and recognized as the “splitting” state until the droplet is totally spitted from the reservoir (Fig. 7d4, Fig. 7c, and †video 4). To create a droplet of one unit size, simply activate one electrode on one side of the reservoir. AI-assisted methods can adjust the ideal size of droplets on demand, and can be used to enhance the precise of droplet generation for high-throughput experiments.

The fundamental droplet operations shown above demonstrate the unprecedented advantages of the AI-assisted feedback control system for DMF. It can autonomously conduct experimental operations with various forms of reagents, such as protein-rich droplets, on DMF chips without human intervention. Moreover, it can combine with reinforcement learning to enhance the precision and automation of droplet manipulation by recognizing, tracking, discriminating states, and automating droplet control, thereby expanding the capabilities and applications of DMF systems.

## 5 Conclusions

In summary, an AI-assisted digital microfluidic system named μDropAI is presented in this paper. The system utilizes the U-net model for multistate droplet recognition and segmentation and employs a closed-loop feedback control strategy for real-time control during multistate droplet manipulations. The U-net-based model is robust enough to recognize droplets at different brightnesses, while it can also segment droplets of different colors and shapes, different states, and even the merging of droplets of different colors. Meanwhile, the AI-assisted DMF platform demonstrates the ability to automatically correct failed droplet manipulations by recognizing the droplet position and judging the manipulation state. We envision that this AI-assisted feedback approach will be widely applied in other DMF platforms to achieve precise and automatic droplet manipulations in a wide range of automated DMF applications. Moreover, with the advancement of large-scale models such as ChatGPT, the AI-assisted digital microfluidic system can be integrated with ChatGPT to enable more intelligent manipulations.

Nonetheless, this study is a preliminary proof-of-concept, and we note several limitations of the current implementation. For one, the model is too large to be integrated for computation on, for example, embedded devices, as this is more computationally time and resource intensive. This is a problem that we plan to start with model reduction to ensure accurate recognition while reducing the number of parameters. In addition, the manipulation of multiple droplets is still a difficult point, and we intend to design algorithms to track the manipulation process of droplets and realize automatic feedback control of the manipulation of multiple droplets.

## Supporting information

Electronic Supplementary Information

## Acknowledgements

This research was funded by the National Key R&D Program of China (2021YFC2101101) and the National Natural Science Foundation of China (No. 62103148).

## References

1. J. Zhong, J. Riordon, T. C. Wu, H. Edwards, A. R. Wheeler, K. Pardee, A. Aspuru-Guzik and D. Sinton, Lab Chip, 2020, 20, 709–716.

2. R. P. S. de Campos, D. G. Rackus, R. Shih, C. Zhao, X. Y. Liu and A. R. Wheeler, Anal. Chem., 2019, 91, 2506–2515.

3. J. Guo, L. Lin, K. Zhao, Y. Song, M. Huang, Z. Zhu, L. Zhou and C. Yang, Lab Chip, 2020, 20, 1577–1585.

4. S. Atabakhsh and S. Jafarabadi Ashtiani, Microfluid. Nanofluid., 2018, 22, 1577–1585.

5. M. Torabinia, P. Asgari, U. S. Dakarapu, J. Jeon and H. Moon, Lab Chip, 2019, 19, 3054–3064.

6. H. Geng, J. Feng, L. M. Stabryla and S. K. Cho, Lab Chip, 2017, 17, 1060–1068.

7. K. Jin, C. Hu, S. Hu, C. Hu, J. Li and H. Ma, Lab Chip, 2021, 21, 2892–2900.

8. L. Wan, T. Chen, J. Gao, C. Dong, A. H. Wong, Y. Jia, P. I. Mak, C. X. Deng and R. P. Martins, Sci. Rep., 2017, 7, 14586.

9. J. X. Wang, J. J. Guo, K. F. Zhao, W. D. Ruan, L. Li, J. J. Ling, R. X. Peng, H. M. Zhang, C. Y. Yang and Z. Zhu, Lab Chip, 2021, 21, 2702–2710.

10. J. Zhai, H. Li, A. H. Wong, C. Dong, S. Yi, Y. Jia, P. I. Mak, C. X. Deng and R. P. Martins, Microsyst. Nanoeng., 2020, 6, 6.

11. Q. Y. Ruan, W. D. Ruan, X. Y. Lin, Y. Wang, F. X. Zou, L. J. Zhou, Z. Zhu and C. Y. Yang, Sci. Adv., 2020, 6, eabd6454.

12. J. Lamanna, E. Y. Scott, H. S. Edwards, M. D. Chamberlain, M. D. M. Dryden, J. Peng, B. Mair, A. Lee, C. Chan, A. A. Sklavounos, A. Heffernan, F. Abbas, C. Lam, M. E. Olson, J. Moffat and A. R. Wheeler, Nat. Commun., 2020, 11, 5632.

13. B. B. Li, E. Y. Scott, M. D. Chamberlain, B. T. V. Duong, S. L. Zhang, S. J. Done and A. R. Wheeler, Sci. Adv., 2020, 6, eaba9589.

14. X. Xu, L. Lin, J. Yang, W. Z. Qian, R. Su, X. X. Guo, L. F. Cai, Z. R. Zhao, J. Song and C. Y. Yang, Nano Today, 2022, 46, 101596.

15. A. A. Sklavounos, C. R. Nemr, S. O. Kelley and A. R. Wheeler, Lab Chip, 2021, 21, 4208–4222.

16. G. Sathyanarayanan, M. Haapala, C. Dixon, A. R. Wheeler and T. M. Sikanen, Adv. Mater. Technol., 2020, 5, 2000451.

17. R. Fobel, C. Fobel and A. R. Wheeler, Appl. Phys. Lett., 2013, 102, 193513.

18. T. S. Kaminski and P. Garstecki, Chem. Soc. Rev., 2017, 46, 6210–6226.

19. M. Abdelgawad and A. R. Wheeler, Adv. Mater., 2009, 21, 920–925.

20. M. J. Jebrail and A. R. Wheeler, Curr. Opin. Chem. Biol., 2010, 14, 574–581.

21. M. J. Jebrail, M. S. Bartsch and K. D. Patel, Lab Chip, 2012, 12, 2452–2463.

22. S. C. C. Shih, R. Fobel, P. Kumar and A. R. Wheeler, Lab Chip, 2011, 11, 535–540.

23. S. Sadeghi, H. J. Ding, G. J. Shah, S. P. Chen, P. Y. Keng, C. J. Kim and R. M. van Dam, Anal. Chem., 2012, 84, 1915–1923.

24. T. Lederer, S. Clara, B. Jakoby and W. Hilber, Microsystem Technologies - Micro- and Nanosystems*. Information Storage and Processing Systems*, 2012, 18, 1163–1180.

25. C. J. Zhang, Y. Su, S. Y. Hu, K. Jin, Y. H. Jie, W. S. Li, A. Nathan and H. B. Ma, ACS Omega, 2020, 5, 5098–5104.

26. J. Gao, X. M. Liu, T. L. Chen, P. I. Mak, Y. G. Du, M. I. Vai, B. C. Lin and R. P. Martins, Lab Chip, 2013, 13, 443–451.

27. J. Gong and C. J. Kim, Lab Chip, 2008, 8, 898–906.

28. P. Q. N. Vo, M. C. Husser, F. Ahmadi, H. Sinha and S. C. C. Shih, Lab Chip, 2017, 17, 3437–3446.

29. Y.-J. Shin and J.-B. Lee, Rev. Sci. Instrum., 2010, 81, 014302.

30. S. Ghosh, H. Rahaman and C. Giri, 2018 IEEE 27th Asian Test Symposium (ATS), Hefei, China, 2018, pp. 185–190.

31. L. B. Li, Z. Gu, J. L. Zhou, B. Y. Yan, C. Kong, H. Wang and H. F. Wang, Chin. Chem. Lett., 2021, 32, 3416–3420.

32. J. Long, E. Shelhamer, T. Darrell and Ieee, Proceedings of the IEEE Conference on Computer Vision and Pattern Recognition (CVPR), Boston, MA, USA, 2015, pp. 3431–3440.

33. M. J. Shafiee, B. Chywl, F. Li and A. Wong, arXiv, 2017, preprint, arXiv:1709.05943.

34. S. A. Taghanaki, K. Abhishek, J. P. Cohen, J. Cohen-Adad and G. Hamarneh, Artificial Intelligence Review, 2021, 54, 137–178.

35. P. Wang, P. Chen, Y. Yuan, D. Liu, Z. Huang, X. Hou and G. Cottrell, 2018 IEEE Winter Conference on Applications of Computer Vision (WACV), Lake Tahoe, NV, USA, 2018, pp. 1451–1460.

36. Y. Mo, Y. Wu, X. Yang, F. Liu and Y. Liao, Neurocomputing, 2022, 493, 626–646.

37. H. Wang, Y. Y. Chen, Y. F. Cai, L. Chen, Y. C. Li, M. A. Sotelo and Z. X. Li, IEEE Transactions On Intelligent Transportation Systems, 2022, 23, 21405–21417.

38. A. Lin, B. Chen, J. Xu, Z. Zhang, G. Lu and D. Zhang, IEEE Trans. Instrum. Meas., 2022, 71, 1–15.

39. L. Wang, J. J. Wu, X. Y. Liu, X. L. Ma and J. Cheng, Complex & Intelligent Systems, 2022, 8, 3833–3845.

40. R. Fobel, C. Fobel and A. R. Wheeler, Appl. Phys. Lett., 2013, 102, 193513.

41. M. Alistar and U. Gaudenz, Bioengineering, 2017, 4, 45.

42. A. Radford, K. Narasimhan, T. Salimans and I. Sutskever, 2018, https://api.semanticscholar.org/CorpusID:49313245.

43. O. Ronneberger, P. Fischer and T. Brox, Medical Image Computing and Computer-Assisted Intervention – MICCAI 2015, Munich, Germany, 2015, pp. 234–241.

44. T. Pavlidis and Y. T. Liow, IEEE Transactions on Pattern Analysis and Machine Intelligence, 1990, 12, 225–233.

45. Z.-R. Song, J. Zeng, J.-L. Zhou, B.-Y. Yan, Z. Gu and H.-F. Wang, Micromachines, 2022, 13, 1563.

46. L.-C. Chen, Y. Zhu, G. Papandreou, F. Schroff and H. Adam, Proceedings of the European Conference on Computer Vision (ECCV), Munich, Germany, 2018, pp. 801–818.

