## Supplementary material for "An Artificial Intelligence-Assisted Digital Microfluidic System for Multistate Droplet Control": Electronic Supplementary Information: ESI.docx

**The electronic supplementary information includes:**

S1. Region growing algorithm;

S2. Equations of the state accuracy, position accuracy, overall accuracy, mean precision and mean pixel accuracy;

S3. The U-net model evaluation index of the pixels;

S4. Recognition and segmentation results under different colours, shapes, sizes, and states;

S5. Droplet splitting and moving and reservoir splitting droplet;

Video 1-4. Experiment results of real-time recognition of the multi-state during droplet manipulations (video S1), and experiment results of automated feedback control of droplet splitting, moving, (video S2 and video S3) and droplet dispensing (video S4).

**S1. Region growing algorithm**

The region growing algorithm is an image segmentation method based on a specific detection criterion, whose basic idea is to form a series of regions by progressively aggregating pixels with similar properties in all directions from an initial point (or a pre-determined seed point) that possesses the ability to satisfy specific conditions. As shown in the Fig. 1, in each segmented region, by selecting a seed pixel as the starting point for growth, subsequently, neighboring pixels that are consistent or similar to that seed pixel under some a priori growth or similarity criterion are fused into the same region. The subsequently fused pixel will be considered as a new seed pixel, which in turn continues the above process. This iterative progression continues until there are no more mergeable pixels that satisfy the growth criteria, thus allowing the region to reach a steady state.

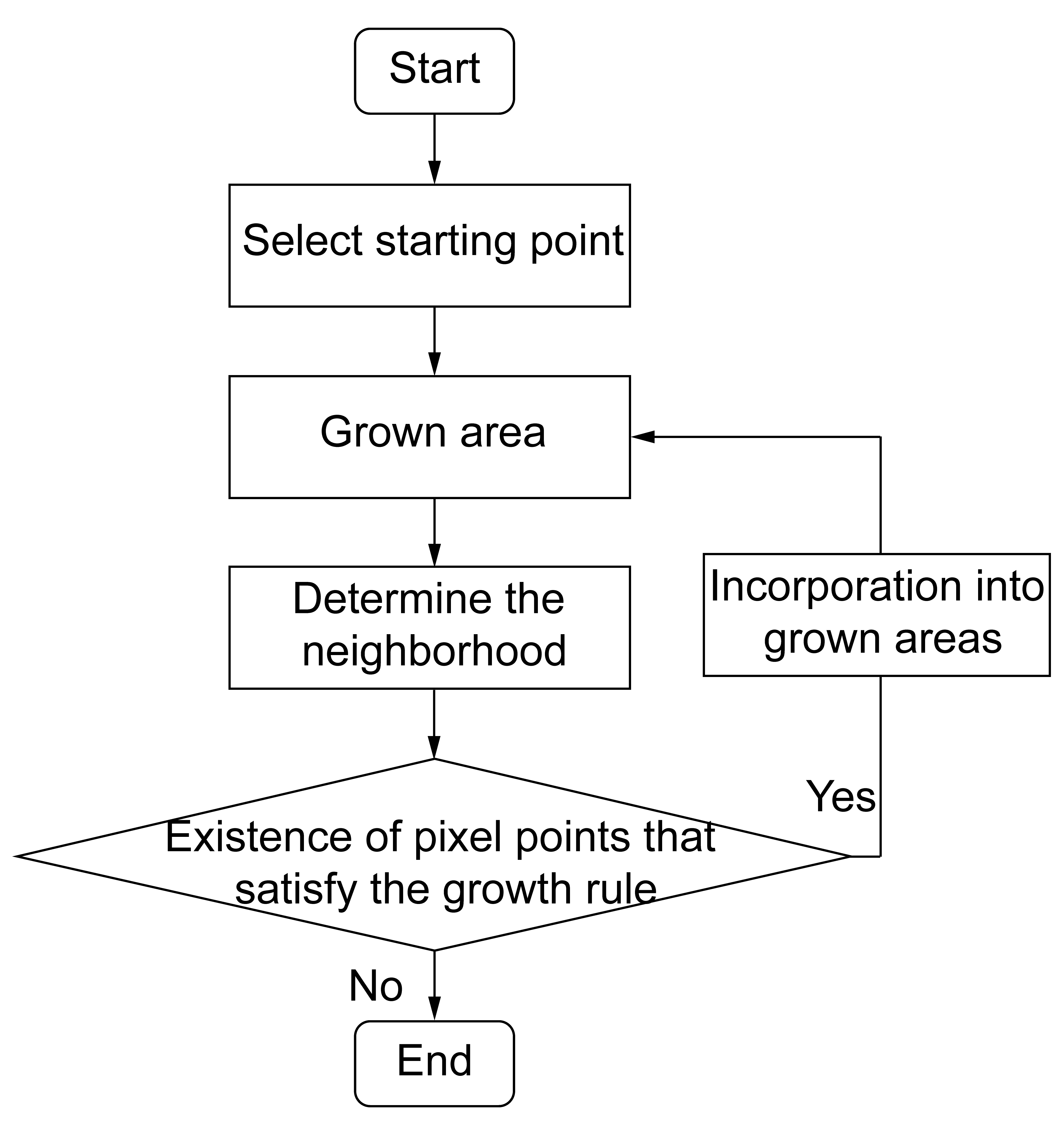

Fig. 1 Flow chart of region growing algorithm

**S2. Equations of the mean precision, mean pixel accuracy, mean recall, state accuracy, position accuracy, and overall accuracy**

The mean precision (mPrecision) refers to a metric that calculates the precision for each category in a multi-class scenario and takes the average of these values. The calculation equation is shown as Equ.1 and Equ.2.

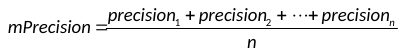

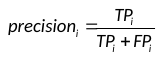

Where,
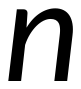
 represents the total number of categories,
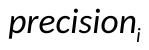
represents the precision of the
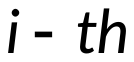
 category.
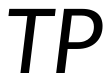
 represents the number of samples correctly predicted as positive by the model in all samples, while
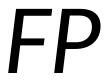
 represents the number of samples incorrectly predicted as positive by the model in all samples.

The mean pixel accuracy (mPA) refers to comparing each pixel predicted by the model with its corresponding ground truth label and calculating the ratio of correctly classified pixels to the total number of pixels, calculated as Equ.3.

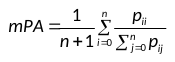

Where,
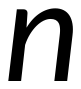
 represents the total number of categories,
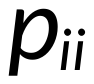
 represents the number of correct predictions,
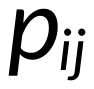
 represents predicting of
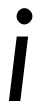
 as
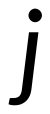
, false negatives.

The Mean Recall (mRecall) is usually the average recall in a multi-category classification task. Used to measure the overall performance of the model in a multi-category classification problem, it considers the recall of each category and averages them. The calculation equation is shown as Equ.4 and Equ.5.

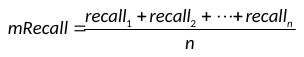

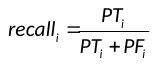

Where,
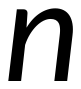
 represents the total number of categories,
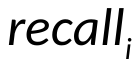
represents the precision of the
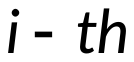
 category.
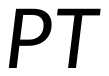
 represents the number of samples correctly predicted as positive by the model in all positive samples, while
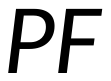
 represents the number of samples incorrectly predicted as positive by the model in all negative samples.

State accuracy is an algorithm's ability to recognize the current state of a droplet in each frame of a video. It is measured by comparing the recognized droplet states with the actual droplet states in the recorded video. The state accuracy is calculated by determining the percentage of correctly recognized droplet states over the total number of frames in the video, as shown in Equ.1.

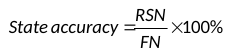

Where,
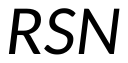
represents the number of frames in which the droplet states are correctly recognized, and
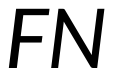
 represents the total number of frames in the real-time video.

Position accuracy refers to the algorithm's ability to recognize the precise position of a droplet in each frame of a video. It is measured by comparing the algorithm's recognized droplet positions with the actual droplet positions in the real-time video. The algorithm determines the droplet's position by identifying the location with the highest concentration of droplet pixels, and this location is considered as the droplet's position. Equ.2 represents the calculation of position accuracy, which is the percentage of correctly recognized droplet positions over the total number of frames in the video.

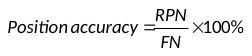

Where,
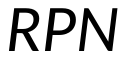
 represents the number of frames in which the droplet positions are correctly recognized, and
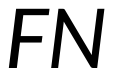
 represents the total number of frames in the real-time video.

Overall accuracy refers to the algorithm's ability to recognize both the state and position of droplets in each frame of a video. It is calculated by comparing the algorithm's recognized droplet states and positions with the actual droplet states and positions in the real-time video. The overall accuracy is determined by calculating the percentage of frames in which both the droplet state and position are correctly recognized, out of the total number of frames in the video, as shown in Equ.3.

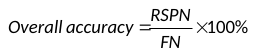

Where,
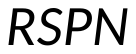
 represents the number of frames in which both the droplet state and position are correctly recognized, and
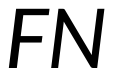
 represents the total number of frames in the real-time video.

**S3. The proposed model evaluation index of the pixels**

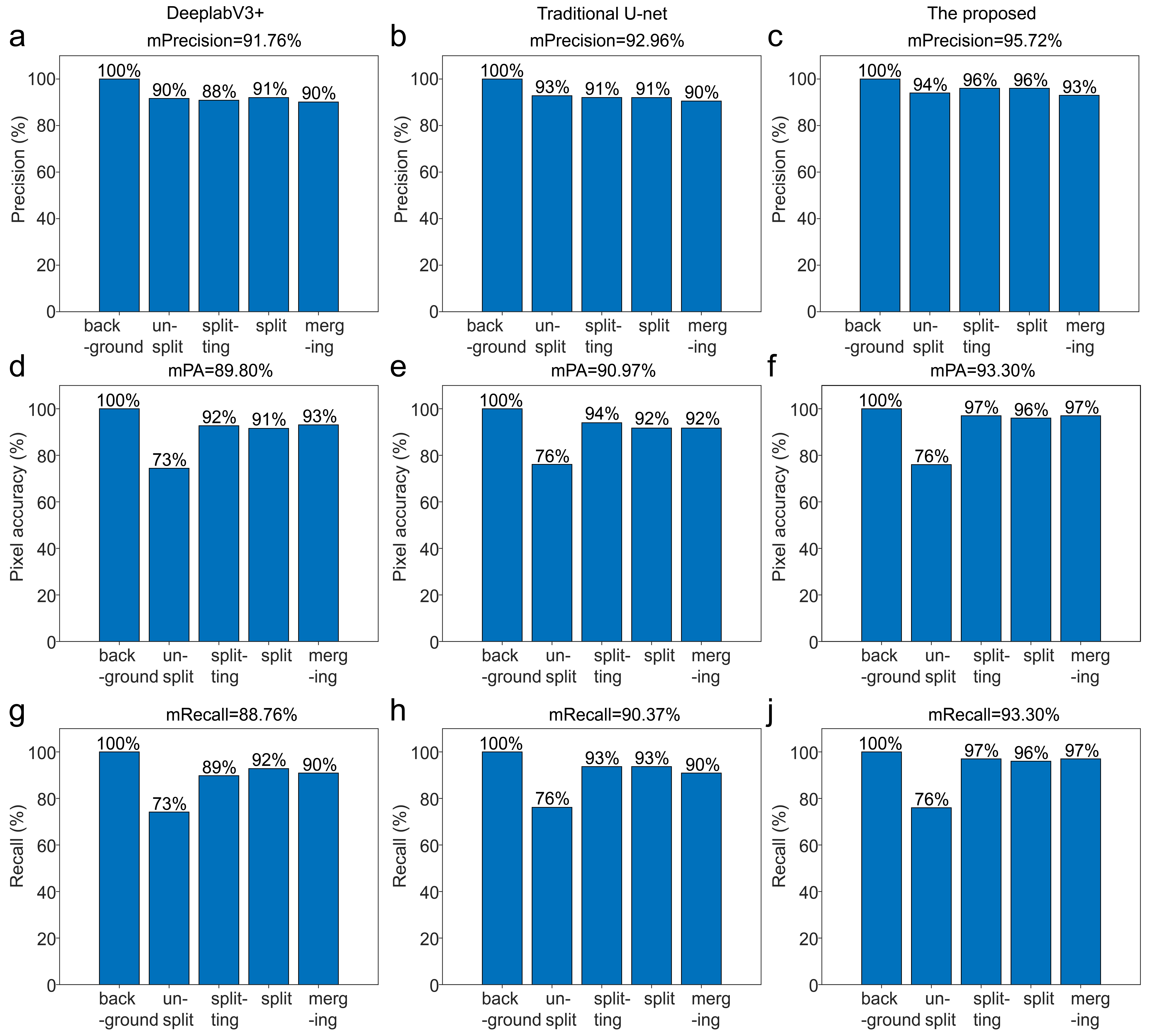

Fig. 2 The comparison evaluation index on the pixels of the DeeplabV3+, traditional U-net, and the proposed. (a) The mPrecision of the DeeplabV3+; (b) The mPrecision of the traditional U-net; (c) The mPrecision of the proposed; (d) The mPA of the DeeplabV3+; (e) The mPA of the traditional U-net; (f) The mPA of the proposed; (g) The mRecall of the DeeplabV3+; (h) The mRecall of the traditional U-net; (j) The mRecall of the proposed.

**S4. Recognition and segmentation results under different colors and shapes**

As shown in Fig. 3, we recognize droplets of different colors and shapes. Fig. 3a shows that the algorithm distinguishes the droplets from the background and accurately recognizes that the droplets are in the "unsplit" state, regardless of whether the droplets are black, yellow, blue, or even transparent, and regardless of whether the droplets are in the shape of a straight line, a square, an l-shape, or a triangle. We also discuss the change in state of droplets of different colors as they divide. When a droplet is hourglass-shaped, it is recognized as a "splitting" state. When a large droplet splits into two small droplets, the splitting process is complete and the state changes to "split". When two separated droplets merge, the state is recognized as "merging" (Fig. 3b-e). The state changes all match the fluidic changes of the droplets during the actual operation, which proves that our system has a strong generalization ability to recognize the instantaneous operational state changes of droplets with different colors and shapes.

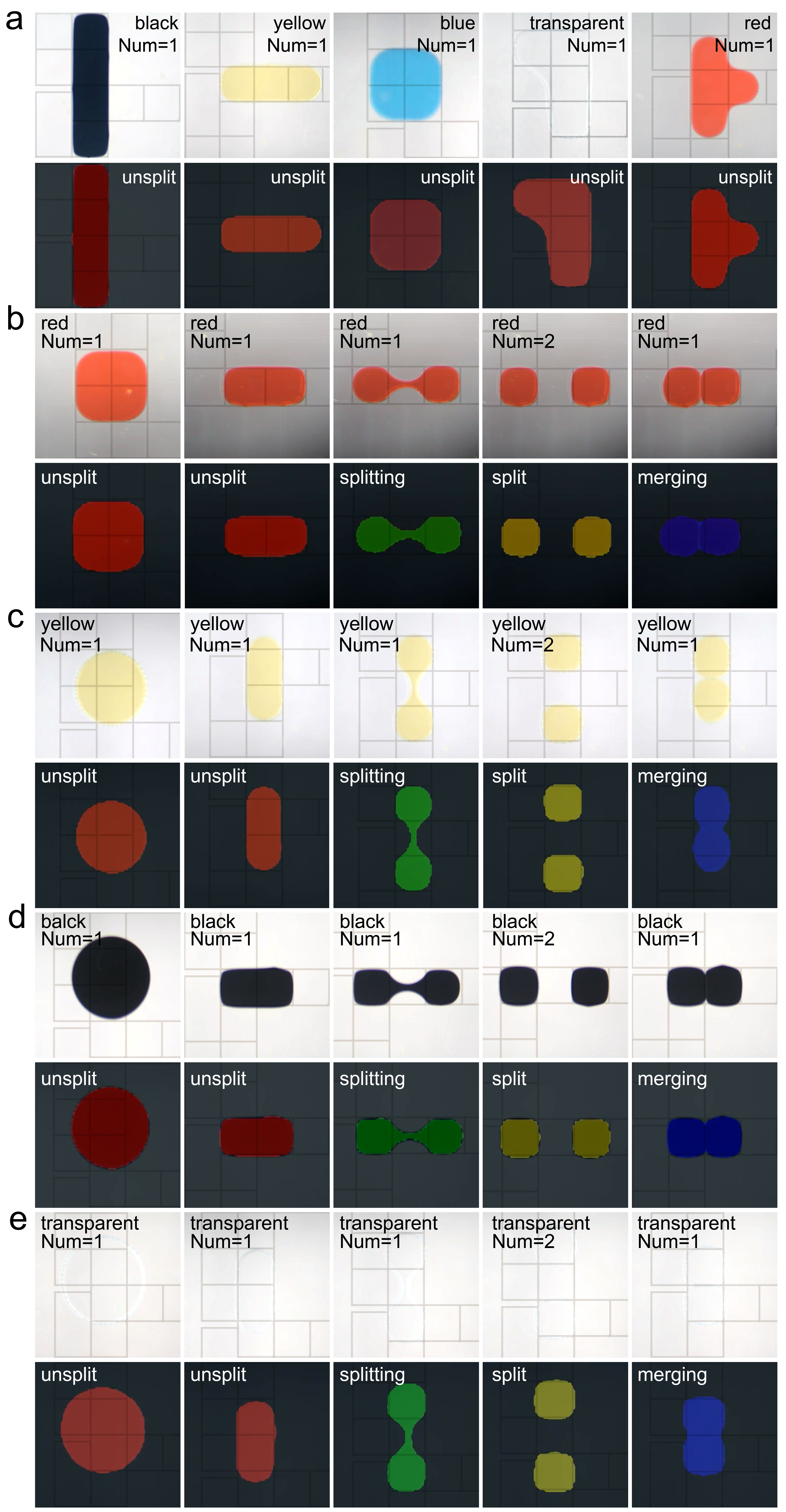

Fig. 3 Recognition and segmentation results under different colors, shapes. (a) The recognition results of different colored and shaped droplets; (b) The recognition states of red droplets; (c) The recognition states of yellow droplets; (d) The recognition states of black droplets; (e) The recognition states of transparent droplets.

**S5. Droplet splitting and moving and droplet dispensing**

A flow chart of automated feedback control involving droplet moving and splitting is performed in the Fig. 4. Droplet splitting: First, we will activate the electrode where the droplet is located, then open the electrodes on both sides of the droplet for stretching, and then close the middle electrode for splitting. The droplet state recognized by the semantic segmentation algorithm is used as a feedback signal to judge whether the droplet splits successfully. When the droplet state is "split", the splitting process is successfully completed. Otherwise, driving the droplet back to the initial state and splitting again.

Droplet movement: First, we will give the next moving position according to the current position of the droplet. After the droplet moves, the droplet position recognized by the semantic segmentation algorithm is used as a feedback signal to judge whether the droplet reaches the specified position. If that fails, the electrodes continue to be sent to the specified position, driving the droplet to move.

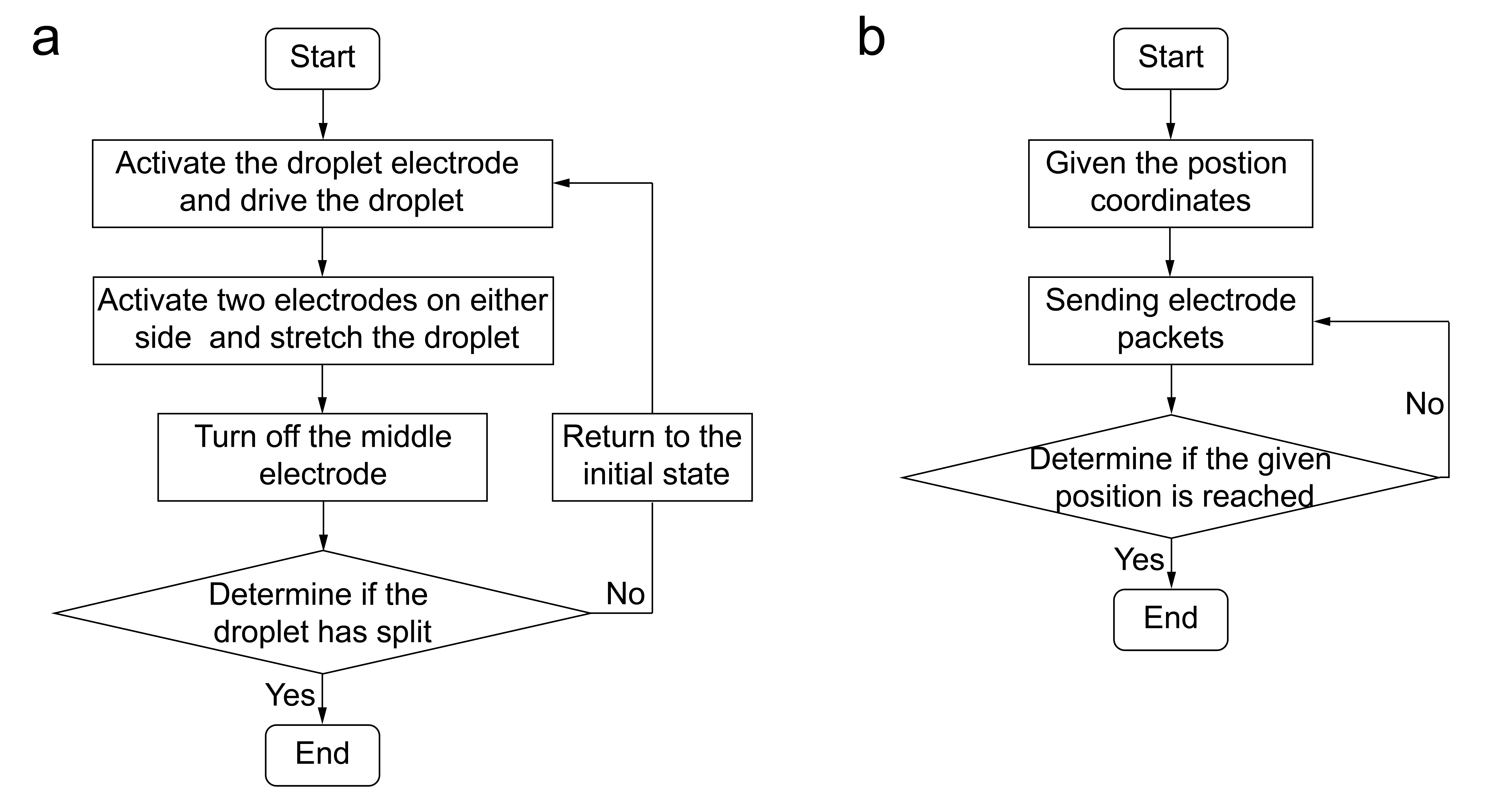

Fig. 4 (a) Flow chart for automated feedback control of droplet splitting; (b) Flowchart of automated feedback control of droplet moving.

A flow chart of automated feedback control experiments for dispensing is performed in the Fig. 5. Droplet dispensing: First, activate the electrode where the reservoir is located. After adding the droplet, open the three electrodes on one side of the reservoir, close the reservoir electrode and stretch the droplet. Activate the reservoir electrode again and close the electrode furthest from the reservoir among the three electrodes to split. Then close the electrode closest to the liquid reservoir, open the remaining two electrodes, and split the droplet. When the droplet state is recognized as "split", the splitting is successful; otherwise, driving the droplet back to the reservoir and operating again.

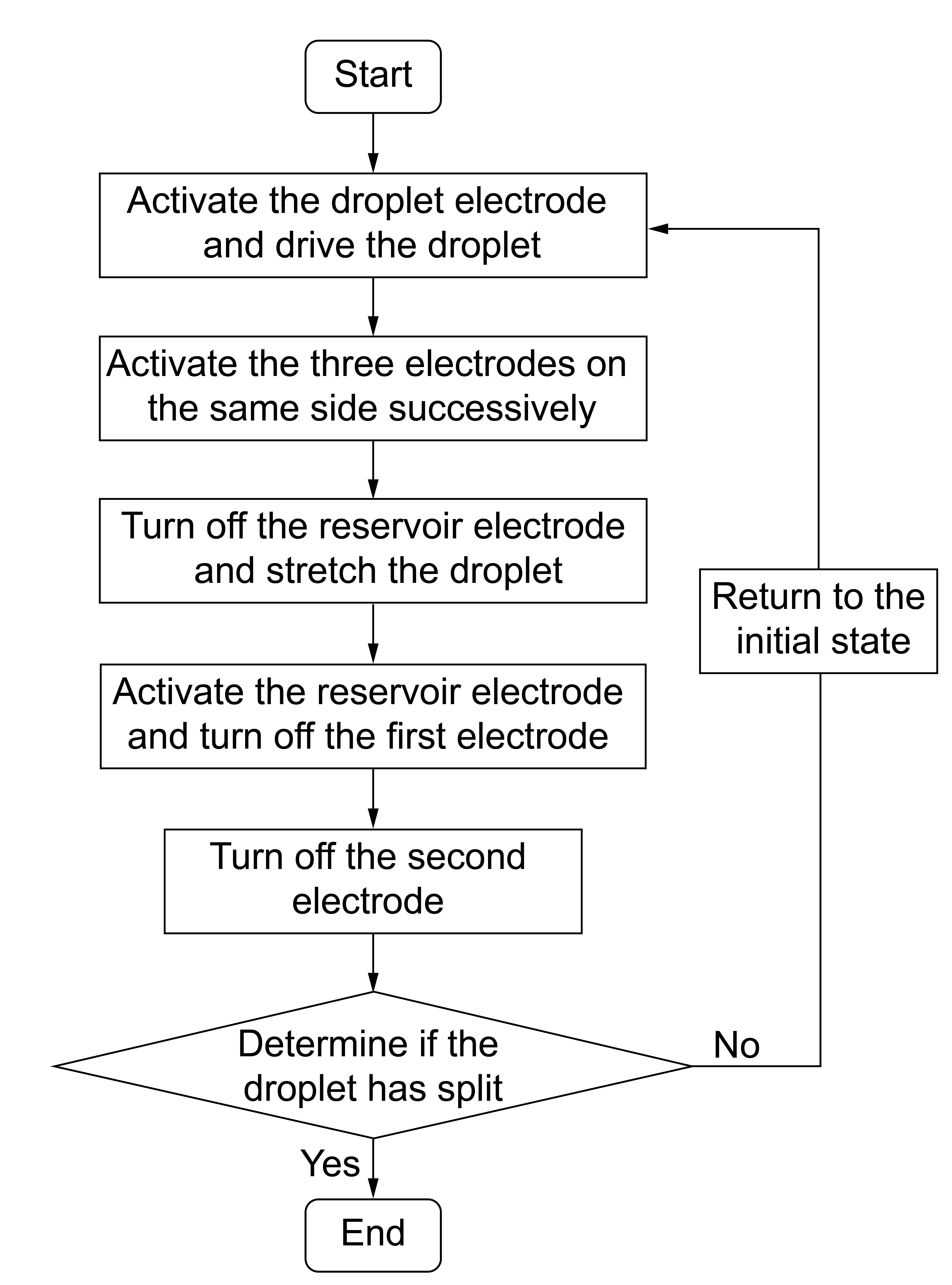

Fig. 5 Flow chart of automated feedback control of droplet dispensing.
